# Validation of scRNA-seq by scRT-ddPCR using the example of *ErbB2* in MCF7 cells

**DOI:** 10.1101/2022.05.31.494164

**Authors:** Tobias Lange, Tobias Groß, Ábris Jeney, Julia Scherzinger, Elly Sinkala, Christoph Niemöller, Stefan Zimmermann, Peter Koltay, Felix von Stetten, Roland Zengerle, Csaba Jeney

## Abstract

Single-cell RNA sequencing (scRNA-seq) can unmask transcriptional heterogeneity facilitating the detection of rare subpopulations at unprecedented resolution. In response to challenges related to coverage and quantity of transcriptome analysis, the lack of unbiased and absolutely quantitative validation methods hampers further improvements. Digital PCR (dPCR) represents such a method as we could show that the inherent partitioning enhances molecular detections by increasing effective mRNA concentrations. We developed a scRT-ddPCR method and validated it using two breast cancer cell lines, MCF7 and BT-474, and bulk methods. *ErbB2*, a low-abundant transcript in MCF7 cells, suffers from dropouts in scRNA-seq and thus calculated fold changes are biased. Using our scRT-ddPCR, we could improve the detection of *ErbB2* and based on the absolute counts obtained we could validate the scRNA-seq fold change. We think this workflow is a valuable addition to the single-cell transcriptomic research toolbox and could even become a new standard in fold change validation because of its reliability, ease of use and increased sensitivity.

## 1 Introduction

RNA-seq is the method of choice for gene expression analysis. Herein, differential expression (DE) analysis between two conditions is pivotal to answer challenging questions in research and clinical applications. In bulk RNA-seq, population heterogeneity remains covert, whereas scRNA-seq can capture delicate differences between cells [1]. The development of new platforms expresses the growing interest in scRNA-seq [2–6] and allows novel applications, such as unmasking transcriptional heterogeneity in healthy and cancerous tissues by functional clustering [7–9], discovering uncharacterized cell types [10], and identifying phylogenetic relationships between cells [11]. scRNA-seq enables researchers to understand underlying mechanisms of drug resistance development and relapse in disease treatment by the detection of rare subpopulations at unprecedented resolution [5,8,12]. However, the inherent low sample input in scRNA-seq introduces a significant amount of noise, which increases the propensity for dropouts and artificially increases cell-to-cell variability [13,14]. This is especially dramatic with respect to low-abundant transcripts, which are often referred to as highly interesting but difficult to reliably analyze [15–18]. Furthermore, the tremendous variety of platforms and bioinformatics tools has not yet solidified into a consistent pipeline [13,19,20]. Additionally, the protocol impacts results, as plate-based Smart-seq2 [21,22] proved to be more sensitive, especially regarding low-abundant transcripts compared to the droplet-based Chromium system from 10X Genomics [23,24].

Thus, DE analysis from scRNA-seq must be independently confirmed by single-cell PCR [25]. Several scRT-qPCR workflows have been described [26–30] as well as a few scRT-ddPCR workflows [31–33]. The majority of these workflows use fluorescence-activated cell sorting (FACS) for single-cell isolation [27,28,30,31,34], while other studies use microfluidic devices [32,33], micromanipulators [26] or manual cell picking [29]. Despite its widespread use, single-cell isolation with FACS requires high sample input and the inherent shear forces can damage the cells and impair RNA integrity [34,35]. Furthermore, qPCR is less sensitive and more susceptible to inhibitors compared to dPCR [17,36], while the detection mechanism of dPCR allows absolute quantification without reference [37]. Particularly, the lower sensitivity of qPCR hampers its use for challenging, single-cell mRNA quantification with a focus on low-abundant transcripts.

Therefore, we here propose a novel method for the validation of fold changes from scRNA-seq. Our scRT-ddPCR method combines gentle (ensuring high cell viability ∼ 80 %) and highly reliable (∼ 90 % single cell isolation efficiency) single cell isolation using the F.SIGHT™ single-cell dispenser (CYTENA GmbH, Freiburg) [38] and contact-free liquid handling (I.DOT, Dispendix, Stuttgart) with highly sensitive dPCR [36]. The F.SIGHT™ requires minimal sample input (down to 5000 cells in 5 µl) and its image-based analysis ensures single-cell isolation and delivers an image proof of each dispensing event, which can be unambiguously assigned to the addressed well of the microplate. Through partitioning, dPCR can reliably detect single molecules and enables absolute quantification [17,32,36]. The latter characteristic is key to inter-experimental comparisons as absolute counts are independent of any standard and depict the ground truth. For the validation of our scRT-ddPCR, we used two breast cancer cell lines, MCF7 and BT-474, the latter overexpresses *ErbB2* [39–41]. We found high concordance between mRNA counts from scRT-ddPCR and bulk RT-ddPCR methods. Interestingly, *ErbB2* log2FCs were significantly different between scRNA-seq and scRT-ddPCR. We assume that the inherent partitioning of dPCR increases sensitivity and resolution, and thus allows us to confirm or reject fold changes from scRNA-seq.

## 2 Materials and methods

### 2.1 Cells and cell culture

MCF7 (ATCC^®^ HTB-22™) and BT-474 (ATCC^®^ HTB-20™) cells were obtained from the BIOSS Centre for Biological Signaling Studies (Freiburg, Germany). MCF7 cells were cultured in DMEM, GlutaMAX™ Supplement (31966021, Gibco™) and BT-474 cells were cultured in DMEM/F12, GlutaMAX™ Supplement (31331028, Gibco™) in Nunc™ EasYFlask™ Cell Culture Flasks (156340, Thermo Scientific™). Both media were supplemented with 10 % FBS (10270106, Gibco™) and 1 % Pen/Strep (15140122, Gibco™). Cells were cultured until ∼90 % conflueny in a cell culture incubator (Heracell™ 150i CO_2_ Incubator, 50116048, Thermo Scientific™) under a 5 % CO_2_ atmosphere at 37 °C. Cells were harvested with 1X TrypLE™ Express Enzyme (12604021, Gibco™). Trypsin activity was quenched by addition of medium. The cells were washed twice with DPBS (14040133, Gibco™) and counted (Countess^®^ II Automated Cell Counter, Invitrogen™) including live/dead staining with trypan blue (T10282, Invitrogen™).

### 2.2 Total RNA isolation and bulk cell lysis

Total RNA was isolated from 1×10^6^ MCF7 and 1×10^6^ BT-474 cells (‘bulk’) using the RNeasy Mini Kit (74104, Qiagen) in combination with the QIAshredder (79654, Qiagen) for lysate homogenization according to manufacturer’s instructions. Simultaneously, total RNA was isolated using the *Quick*-DNA/RNA Microprep Plus Kit (D7005, Zymo Research) with an upfront proteinase K digest and on-column DNase I digest. RNA concentration (**Tab S3**) was measured with the NanoDrop™ One (Thermo Scientific™). RNA integrity was checked on a 1.2 % native agarose gel (2267.1, Roth) using 1X TBE buffer (3061.1, Roth) (**Fig S2c**). 1 µg total RNA was combined with 1X DNA Orange Loading Dye (R0631, Thermo Scientific™) and 60 to 75 % formamide (6749.3, Roth) and heated to 65 °C for 5 min before loading. RNA was visualized with 1X GelRed® Nucleic Acid Stain (SCT123, Milipore). Total RNA was diluted 1:20, 1:50, 1:100 and 1:1000 with PBS for MCF7 cells and 1:50, 1:100, 1:1000 and 1:10000 with PBS for BT-474 cells. Each sample of the dilution series was analyzed regarding *ErbB2* and *ACTB* mRNA counts in triplicates using dPCR (2.5 Droplet digital PCR). The absolute gene mRNA counts per single cell were calculated by dividing the detected number of mRNAs with the number of cells (with respect to the dilution factor). Crude lysates (‘cl’) from 1×10^6^ MCF7 and 1×10^6^ BT-474 cells were prepared using 500 µl LBTW (lysis buffer from PICO Amplification Core (AMC) Kit, PICO-000010, Actome) proprietary buffer of Actome GmbH). The samples were incubated on ice for 5 min, sonicated for 1 min and cell debris were removed by centrifugation at 14000 xg at 4 °C for 10 min. The lysate was diluted with 49.5 ml DPBS (100X dilution), resulting in 20 cell equivalents per µl. Thus, dispensation of 50 nl in to the ddPCR master mix using the I.DOT (2.3 Liquid dispensation using I.DOT) resulted in an equivalent amount of material to a single cell.

### 2.3 Liquid dispensation using I.DOT

The Immediate Drop-on-demand Technology (I.DOT One; Dispendix, Stuttgart, Germany) [42,43] with I.DOT PURE plates 90 µm orifice (Dispendix, Stuttgart, Germany) was used to dispense 0.5 µl LBTW into a 384-well V-bottom plate (0030623304, Eppendorf^®^) for single-cell dispensation (2.4 Single-cell dispensation using F.SIGHT™). The reduced volumes for down-scaled SMART-Seq® (2.6.1 cDNA synthesis using SMART-Seq® Single Cell Kit) and down-scaled library preparation (2.6.2 Library preparation using Nextera XT and sequencing) were dispensed using the I.DOT. Prior to dispensation, the I.DOT was calibrated for the applied liquids to ensure reliable dispensation.

### 2.4 Single-cell dispensation using F.SIGHT™

The single cell dispensing procedure was performed as described earlier [38,44,45]. The F.SIGHT™ single-cell dispenser (CYTENA GmbH, Freiburg, Germany) is an improved version of the single-cell printer (SCP) [38]. Both MCF7 and BT-474 cell concentrations were adjusted to 1×10^6^ cells/ml and loaded into a Dispensing Cartridge (CYTENA GmbH, Freiburg, Germany). The settings for MCF7 cells were 10 to 25 µm cells size (BT-474: 10 to 30 µm) and 0.5 to 1 roundness (same for BT-474) (**Fig 1a, S3a** and **S3b**). The F.SIGHT™ can reliably dispense single cells in minimal liquid volumes [45] within a short period of time (96 single cells per approx. 10 min). The single-cell dispensation efficiency (fraction of successful single-cell isolation events from targeted single-cell isolation events) is usually around 90 % [38], and is additionally controlled by cell images unambiguously assigned to each dispensation event (**Fig 1a**). Thus, other than single-cell dispensation events like multiple cells per droplet or empty droplets can be excluded.

**Figure 1:**
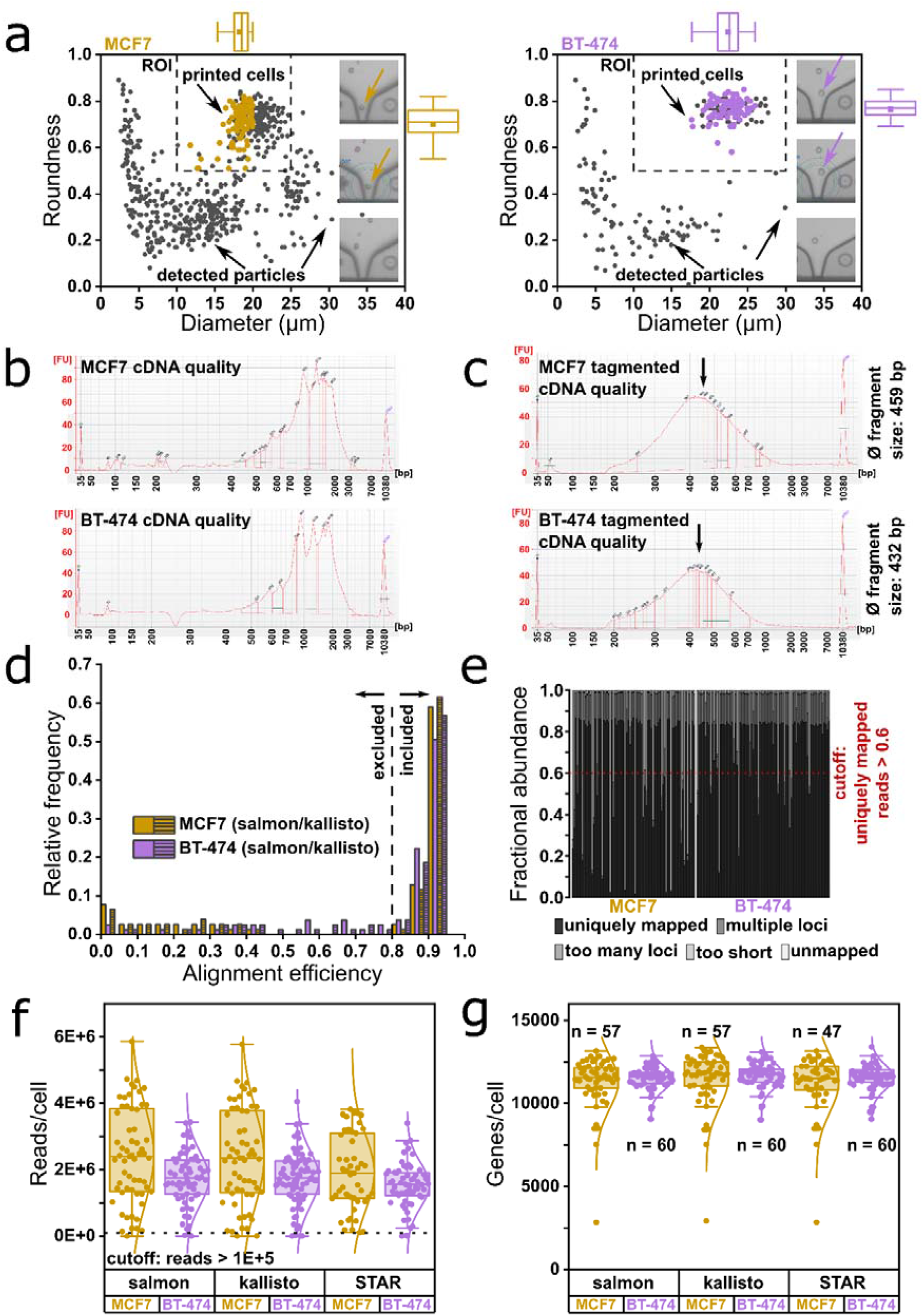
Quality control of down-scaled SMART-Seq® workflow with MCF7 and BT-474 cells. **a)** 2D-scatter plots (roundness vs. diameter) of detected particles in the dispensation nozzle during the process and dispensed cells (colored dots). The particles can be of various origins: cell debris, cell aggregates, corpuscular materials from the cell culture medium or cells. The ROI (region of interest) depicts the desired morphological criteria by which a particle is defined as a cell. The overlap in the ROI between detected particles and dispensed cells is because of the fact that some cells could not be isolated. Boxplots show roundness and diameter distributions of dispensed single cells (n = 84). Representative images of the printing process, which enable manual image-based exclusion of droplets with multiple cells or empty droplets are shown. These images can be unambiguously assigned to the addressed wells of the microplate. **b, c)** Representative electropherograms (Agilent’s Bioanalyzer) of cDNA and tagmented cDNA size distributions for both cell lines. The average cDNA length after tagmentation was 459 bp for MCF7 and 432 bp for BT-474 cells. **d)** Alignment efficiency of salmon and kallisto aligner. Cells with less than 80 % alignment efficiency were excluded from further analyses. **e)** Alignment statistics (fraction of uniquely mapped reads, fraction of reads mapped to multiple loci, fraction of reads mapped to too many loci, fraction of reads too short for mapping, fraction of unmapped reads) for MCF7 and BT-474 cells using STAR aligner. Cells with less than 60 % of uniquely mapped reads were excluded from further analyses. **f)** Total read counts per cell for MCF7 and BT-474 cells using salmon, kallisto or STAR aligner. Cells with less than 1E+5 transcripts were excluded from further analyses. **g)** Gene per cell counts for MCF7 cells (median across all aligners: 11676 genes per cell) and BT-474 cells (median across all aligners: 11682 genes per cell) after all steps of filtering. The median of genes per cell is the same independent of aligner and cell line (p > 0.05, Mann-Whitney test with Bonferroni correction). The amount of cells excluded after each filtering step is shown in **Tab S1**.

### 2.5 Droplet digital PCR

*ErbB2* and *ACTB* mRNAs were analyzed using the naica^®^ Crystal Digital PCR System (Stilla Technologies, Villejuif, France) [46]. Master mix was prepared as follows: 11.5 µl qScript XLT 1-Step RT-qPCR ToughMix (2X) (95132, Quantabio), 1.15 µl TaqMan Assay Hs01001580_m1 (20X) (ErbB2) (4331182, Applied Biosystems™), 1.15 µl TaqMan Assay Hs01060665_g1 (20X) (ACTB) (4448489, Applied Biosystems™), 0.23 µl fluorescein (100X, prepared according to “Fluorescein preparation for naica^®^ system” from Stilla Technologies) (0681-100G, VWR Chemicals), 1 µl of diluted RNA sample (or a single cell), ad 23 µl H_2_O. After thorough mixing, 20 µl of the reaction mix were transferred bubble-free to the chambers of the Sapphire Chips (Stilla Technologies, Villejuif, France). The dPCR conditions of the Geode cycler were: partitioning of the reaction mix, cDNA synthesis (50 °C, 10 min), initial denaturation (95 °C, 1 min); followed by 45 cycles of denaturation (95 °C, 6 s), annealing and extension (60 °C, 45 s) and finally the pressure was released. The chips were transferred to the Prism3 reader and imaged using exposure times: 65 ms and 150 ms for FAM and HEX channel (82 mm focus). Afterwards, droplet quality was manually controlled and in case of poor quality, e.g. coalescence or air bubbles, the respective areas were excluded from further analysis. All NTCs were negative (data not shown). The average droplet volume using 1X qScript XLT 1-Step RT-qPCR ToughMix is 0.548 nl. Hence, the corresponding analysis configuration file was used for quantification (User Manual v2.1 of the Crystal Miner Software, Stilla Technologies, 2018). We comply with the dMIQE guidelines [47,48] and report all essential information (**Tab S7**).

### 2.6 Down-scaled single-cell RNA sequencing

#### 2.6.1 cDNA synthesis using SMART-Seq® Single Cell Kit

For cDNA synthesis the SMART-Seq® Single Cell Kit (634472, Takara BIO) at 1/10 of the original reaction volume was used (down-scaled SMART-Seq® workflow). Briefly, 1.15 µl of the lysis buffer (0.1 µl Reaction Buffer, 0.1 µl 3’ SMART-Seq CDS Primer II A and 0.95 µl dH_2_O) were dispensed in skirted 384-well PCR plates (4ti-0384/X, 4titude) using the I.DOT. Subsequently, single cells (84 cells per cell line) and NTCs (empty droplets) were dispensed into the lysis buffer. The plates were sealed (AB0558, Thermo Scientific™) and snap-frozen at -80 °C until further processing. After thawing, primers were annealed in a C1000 Touch™ Thermal Cycler (Bio-Rad Laboratories, Hercules, CA, USA) at 72 °C for 3 min. Afterwards, 0.75 µl of RT Master Mix (0.4 µl SMART-Seq sc First Strand Buffer, 0.1 µl SMART-Seq sc TSO, 0.05 µl RNase Inhibitor and 0.2 µl SMARTScribe II Reverse Transcriptase) were added to each well using the I.DOT. The plate was processed in the C1000 Thermal Cycler for cDNA synthesis at 42 °C for 180 min, 70 °C for 10 min and 4 °C hold. Next, 3 µl PCR Master Mix (2.5 µl SeqAmpCB PCR Buffer (2X), 0.1 µl PCR Primer, 0.1 µl SeqAmp DNA Polymerase and 0.3 µl dH_2_O) were added to each well and cDNA was amplified (95 °C for 1 min, 19 cycles: 98 °C for 10 sec, 65 °C for 30 sec, and 68 °C for 3 min; 72 °C for 10 min and 4 °C hold). Purification of cDNA was performed manually using 9 µl of AMPure XP bead suspension (A63880, Beckman Coulter) per well according to manufacturer’s instructions. In brief, beads and cDNA were incubated for 8 min at room temperature. The beads were separated using conventional neodym magnets for 5 min and beads were washed with 30 µl 80 % ethanol for 30 sec. Afterwards, the beads were resuspended in 17 µl 10 mM Tris-HCl and incubated for 8 min at room temperature. The beads were separated using magnetic separation for 5 min. 15 µl supernatant of each well were transferred to a fresh 384-well plate. The cDNA quantity was determined with the Quant-iT™ PicoGreen™ dsDNA Assay Kit (P7589, Invitrogen™) in 384-well plates (4ti-0203, 4titude). Fluorescent intensities were measured using the Spark 10M Microplate Reader (Tecan, Männedorf, Switzerland). cDNA quality was determined with Agilent’s 2100 Bioanalyzer using the High Sensitivity DNA Kit (5067-4626, Agilent) according to manufacturer’s instructions (representative images in **Fig 1b**).

#### 2.6.2 Library preparation using Nextera XT and sequencing

Prior to tagmentation cDNA was normalized to 0.2 ng/µl. Tagmentation was performed using the Nextera XT DNA Library Preparation Kit (FC-131-1024, Illumina) at 10-fold down-scaled reaction volumes. 1 µl of Tagment DNA Buffer and 0.5 µl Amplicon Tagment Mix were added to each well using the I.DOT. The amplified and purified cDNA was tagmented for 8 min at 55 °C in a C1000 Thermal Cycler. Transposase activity was quenched by addition of 0.5 µl Neutralize Tagment Buffer using the I.DOT (< 1 min) and incubation at room temperature for 5 min. MCF7 and BT-474 libraries were independently amplified with index primers N7xx and S5xx of Nextera XT Index Kit v2 Set A (FC-131-2001, Illumina) and Nextera XT Index Kit v2 Set B (FC-131-2002, Illumina). 1.5 µl Nextera PCR Master Mix and 0.5 µl of a unique combination of primers were added to each well using the I.DOT and tagmented cDNA was amplified in a C1000 Thermal Cycler (72 °C for 3 min, 95 °C for 30 sec, 12 cycles: 95 °C for 10 sec, 55 °C for 30 sec, and 72 °C for 30 sec; 72 °C for 5 min and 10 °C hold). The cDNA libraries for MCF7 and BT-474 were pooled separately (total volume ∼ 420 µl), and purified according to the previously mentioned bead clean-up procedure (using 0.6 to 1-fold AMPure XP bead suspension, A63880, Beckman Coulter), except that after removal of ethanol, 75 µl of resuspension buffer were added and incubated for 3 min. 73 µl of eluted library were transferred to a fresh tube. The quality of tagmented libraries was determined using the High Sensitivity DNA Kit on Agilent’s 2100 Bioanalyzer (representative images in **Fig 1c**). The pooled library was sequenced on the NextSeq 500 System (Illumina, San Diego, CA, USA) using the High-Output v2.5 Kit (20024906, Illumina) with 75 bp single-end reads.

#### 2.6.3 Bioinformatics data analysis

FASTQ files were generated with bcl2fastq v2.20 (“--no-lane-splitting” flag). Sample quality was assessed with FASTQC v0.11.9 (exemplary images: **Fig S1**). The aligners, salmon v1.3.0 [49], kallisto v0.46.1 [50] and STAR 2.7.5c [51], were wrapped into bash scripts and the FASTQ files were separately aligned aligned to the GRCh38 cDNA reference transcriptome from Ensembl using salmon (“--validateMappings” flag) or kallisto as well as to the GRCh38.p13 genome with STAR in solo mode. As recommended for salmon and kallisto, the mean read length and the standard deviation were calculated for each file. For STAR aligner, a genome index was calculated. The outputs were analyzed in Jupyter Notebooks [52] (jupyter core v4.7.0, jupyter-notebook v6.1.6) using R version 4.1.2. Transcript abundances (TPM) and count estimates were imported with the tximeta [53] package for salmon and tximport [54] for kallisto and summarized to genes using the summarizeToGene() statement. STAR alignment files were counted with featureCounts v2.0.3 [55] and the count matrices were directly imported. Cells with an alignment efficiency below 80 % were filtered out (salmon and kallisto) (**Fig 1d** and **Tab S1**). Cells with less than 60 % of uniquely mapped reads were filtered out (STAR) (**Fig 1e** and **Tab S1**). In both cases cells with at least 1E+5 detected reads were considered. Pseudo-bulk differential expression analysis was performed with DESeq2 v1.32.0 [56] on count matrices using LRT testing and suggested parameters for single-cell testing. In order to evaluate transcriptional similarity between cells assayed using our down-scaled SMART-Seq® (salmon aligner only) and published data (GSE151334) [4], the datasets were concatenated into a single AnnData object and imported into SCANPY (v1.8.1) [57]. Cells with fewer than 200 genes expressed and genes expressed in less than three cells were excluded from further analysis. Counts per cell were normalized with SCANPY’s built-in normalization method and log-transformed according to the standard workflow recommended in the SCANPY documentation. Batch-correction was performed with BBKNN [58]. Dimensionality reduction was performed with SCANPY’s built-in UMAP-function (uniform manifold approximation and projection). Scripts for the here described analysis are available from github.com/LangeTo/scRNA-seq_scripts.

### 2.7 Statistical analysis

Groups were initially tested for normal distribution (Shapiro-Wilk test) and for homoscedasticity (F-test) upon which information a suitable test was chosen (if not stated differently): Mann-Whitney test (normal distribution rejected), Welch’s t-test (normal distribution, heteroscedasticity) or Student’s t-test (normal distribution, homoscedasticity). Distributions were compared using the Kolmogorov-Smirnov test. Bonferroni correction was applied for multiple testing corrections. Significance levels are indicated as follows: *** p < 0.001, ** p < 0.01, * p < 0.05, ns p > 0.05. Boxplots indicate the inner quartiles of the data (25 % to 75 %). Whiskers show 1.5xIQR (interquartile range). The median is drawn as a horizontal line. The mean is represented by a square. Individual data points are shown as dots. All plots and statistical analyses were performed with OriginPro 2021 (OriginLab Corporation).

### 2.8 Bootstrapping comparison

To compare fold changes between two methods (*a*: scRNA-seq and *b*: scRT-ddPCR), we used a bootstrapping comparison because regular statistical tests suffer from p-value inflation after repetitive bootstrapping. The algorithm is described in **Fig S4** based on the ratio *r*_*g*_ (**Eq 1**). In brief, four arrays of expression values are needed (one per cell line and per method). A subset of each initial expression array is randomly subsampled with replacement (same length as initial array). The log2FCs of the means of these expression arrays are calculated per method. Then the ratio *r*_*g*_ of these log2FCs is determined (**Eq 1**). Subsampling and ratio calculation is repeated 1,000 times. Finally, mean and 95 % confidence intervals (CI) of the new array *r*_*g*_ is determined. If the 95 % CI overlaps with 1, the methods are assumed to yield the same fold change.

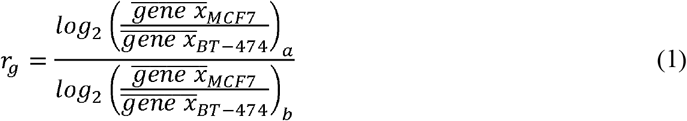

## 3 Results

### 3.1 Down-scaling of SMART-Seq®

It is hypothesized that down-scaling of reaction volumes improves sensitivity [59], while conserving data quality [4,60–62]. This idea follows the concept of dPCR [36], where down-scaling (by partitioning) is an inherent feature, which enhances molecular detections by increasing the effective concentration of nucleic acids. Thus, we down-scaled our scRNA-seq reaction volumes to yield the most precise log2FCs. We used the F.SIGHT™ (CYTENA GmbH, Freiburg) for single cell isolation and the I.DOT (Dispendix, Stuttgart) for contact-free liquid handling. The F.SIGHT™ uses a microfluidic chip generating free-flying, picoliter-sized droplets in which single cells are encapsulated and delivered to the microplate [34,35,38]. Image-based analysis intercepts a permanent vacuum suction when single cells of the desired morphological criteria are detected in the nozzle. High precision is ensured by automatic dispenser offset compensation (AOC) enabling single-cell deposition into few hundred nanoliters in 384-well plates [45]. Simultaneously, the F.SIGHT™ records an image series for each dispensation event, which can be unambiguously assigned to the addressed well of the microplate [38]. Based on the images, the cells can be qualitatively stratified according to roundness and size from a heterogeneous population of particles (**Fig 1a**, only colored dots are dispensed cells, the grey dots are either artefacts or cells that could not be isolated) resulting in a homogeneous cell population (**Fig 1a**, boxplots at the edges). We manually analyzed all images from putative single cells (84 cells per cell line) and found that 7 % (MCF7) and 4 % (BT-474) were doublets or empty droplets (**Tab S1**). The cells were directly dispensed into the lysis buffer and processed by down-scaled SMART-Seq® and down-scaled Nextera XT protocols (2.6.1 cDNA synthesis using SMART-Seq® Single Cell Kit and 2.6.2 Library preparation using Nextera XT and sequencing). The average fragment length for tagmented cDNA was 459 bp and 432 bp for MCF7 and BT-474 cells, respectively. According to Jaeger *et al*. [62], this is an indication for good quality, tagmented cDNA. Representative electropherograms of cDNA and tagmented cDNA are shown in **Fig 1b** and **1c**. FastQC analysis revealed an average Phred score of above 30 for both cell lines (**Fig S1**). We analyzed sequencing data using three common aligners: salmon [49], kallisto [50] and STAR [51]. Based on alignment efficiency ≥ 80 % (salmon and kallisto) or fraction of uniquely mapped reads ≥ 60 % (STAR), cells of poor quality were excluded from downstream analysis (**Fig 1d** and **1e** and **Tab S1**). Further on, cells with less than 1E+5 reads were also excluded from analysis (**Fig 1f** and **Tab S1**). All aligners yielded the same number of genes per cell. 11676 genes per single MCF7 cell and 11682 genes per single BT-474 cell were detected (**Fig 1g**). We also clustered our data with external data from Isakova *et al*. [4], who used 10-fold down-scaled Smart-seq2 protocol. The clusters of MCF7 cells exactly overlap, while other cells formed independent clusters like the reference cell lines HEK293T and fibroblasts (**Fig 2a**). We performed pseudo-bulk DE analysis with different input to DESeq2 (salmon, kallisto or STAR aligner count matrices) (**Fig 2c, 2d** and **Tab S2**). Interestingly, salmon and kallisto predict the highly significant overexpression of *OLFML3, RAMP3* and *VWA5A* (only salmon), which we could not observe with STAR aligner (**Fig 2c**). To our knowledge there is no supporting evidence for this overexpression in MCF7 cells in literature. Furthermore, STAR aligner input to DESeq2 predicts clearly more DEGs in MCF7 than the other two aligners (**Fig 2d**). We found that *ErbB2* is significantly overexpressed in BT-474 cells as previously described [39], while *ACTB* as a housekeeping gene is not significantly different between the cell lines (**Fig 2c** and **Tab S2**). These findings are consistent across all aligners. Additionally, we evaluated the expression of two marker genes for MCF7 cells, *KRT8* and *TFF1* [4]. *TFF1* is overexpressed in MCF7 cells, while *KRT8* shows differential expression only with STAR aligner input to DESeq2 (**Fig 2c** and **Tab S2**). This underlines furthermore the dissimilarity of the aligners used.

**Figure 2:**
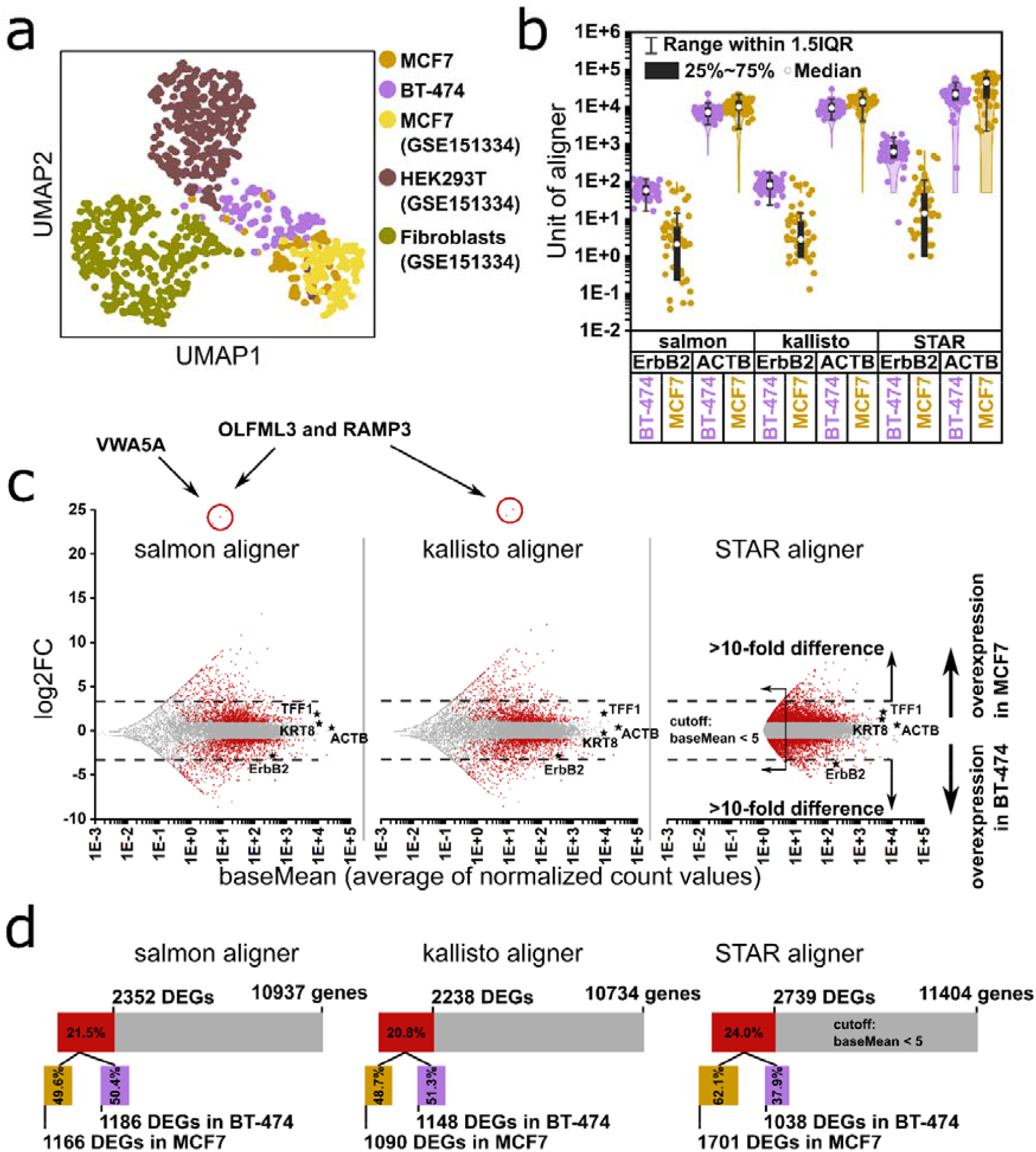
Validation of down-scaled Smart-seq2 by clustering and different bioinformatics pipelines. **a)** UMAP clustering of MCF7 and BT-474 cells along with MCF7 cells, HEK293T cells and fibroblasts from GSE151334. **b)** Violin plots of *ErbB2* and *ACTB* expression values in MCF7 and BT-474 cells from salmon, kallisto and STAR aligner in the respective units (salmon and kallisto: TPM: transcripts per kilobase million; STAR: raw counts). **c)** Bland-Altman plots of gene expression in MCF7 over BT-474 cells with salmon, kallisto or STAR aligner input to DESeq2. Each dot symbolizes a gene with its average expression value in both cell lines (baseMean; x-axis) and the log2FC (log2 of fold change; y-axis). Dots colored in red are significantly differentially expressed genes (DEGs); log2FC > 1 and p_adj_ < 0.05: overexpression in MCF7 cells; log2FC < -1 and p_adj_ < 0.05: overexpression in BT-474 cells. *ErbB2, ACTB, TFF1* and *KRT8* expression values are highlighted with stars and extreme values are highlighted with a red circle. The dashed line indicates 10-fold overexpression in either cell line. **d)** Total number of genes analyzed by DESeq2 using salmon, kallisto or STAR aligner input (STAR: baseMean > 5 necessary for consideration) with amount of DEGs overexpressed in either cell line.

### 3.2 Validation of scRT-ddPCR using bulk methods

For scRT-ddPCR, we isolated cells on the same day and from the same culture as in the case of down-scaled SMART-Seq® using F.SIGTH™ and I.DOT except that the cells were dispensed into LBTW (**Fig S3a** and **S3b**). After lysis, the gene mRNA per cell counts were determined directly form the lysate using digital PCR. First, we investigated varied volumes of LBTW lysis buffer, as the carry-over of detergents may impair droplet formation or reverse transcription and thus PCR efficiency [31,63]. The results indicate that as the volume of lysis buffer increases, the number of formed droplets decreases (**Fig 3a**) due to increased areas of coalescence (**Fig S2a**). The use of 0.5 µl LBTW produces no areas of coalescence and the number of droplets is not significantly reduced, despite a drop of ∼18 % in total droplet number (**Fig 3a**). Similarly, we could not detected a significant difference between the *ErbB2* and *ACTB* mRNA concentrations upon different volumes of lysis buffer (**Fig S2b**). Thus, we used 0.5 µl lysis buffer in subsequent experiments. Of note, at low target concentrations the subsampling error becomes significant [36] due to the loss of mRNAs in the non-partitioned (dead) volume (∼34 % according to manufacturer’s information). We reduced the loaded master mix volume without performance effects (data not shown) to minimize the loss of transcripts (∼18 % dead volume). The well-documented differential expression of *ErbB2* in MCF7 and BT-474 cells [39–41] was taken advantage to demonstrate the ability of our scRT-ddPCR for absolute quantification. We could observe an expression of 9 *ErbB2* mRNA/MCF7 cell (79 % CV), while BT-474 cells expressed a ∼50-fold higher amount (453 *ErbB2* mRNA/BT-474 cell (42 % CV)) (**Fig 3b** and **Tab S5**). Durst *et al*. could observe the same fold difference [39]. On the other hand, we could not detect a significant difference in *ACTB* expression between the cell lines (66 *ACTB* mRNA/MCF7 cell with 44 % CV and 114 *ACTB* mRNA/BT-474 cell with 70 % CV, p > 0.05, Mann-Whitney test with Bonferroni correction; **Fig 3b** and **Tab S5**). The F.SIGHT™ records morphological details of each dispensed cell but we could not detect any correlation between cell size and number of mRNAs per singe cell (**Fig S3c** and **S3d**). These mRNA counts could be biased by incomplete lysis of the single cell. To verify these mRNA counts, we checked the ability of the lysis buffer (LBTW) to exert full dispersion of cell material prior to compartmentalization. Thus, we used two commercially available methods for total RNA isolation, for which we assume a 100 % isolation efficiency (‘bulk’). The two methods differ in sample preparation (DNase I digest vs. no digest and enzymatic lysate homogenization vs. mechanical lysate homogenization), buffers and handling in general, but resulted in the same amount of *ErbB2* or *ACTB* mRNAs per cell (**Fig S2d**). Of note, the RNA quality differs between the two methods (**Fig S2c** and **Tab S3**). Similarly, a single-cell volume equivalent from a crude lysate (‘cl’) cells was dispensed into the dPCR mix and *ErbB2* and *ACTB* counts per single cell were determined to validate that the lysis conditions have no effect on dPCR or the detection of the transcripts. Comparing these extraction methods (‘sc’, ‘bulk’, ‘cl’) for BT-474 cells yields no significant differences for both genes *ErbB2* or *ACTB* (**Fig 3b**). Also *ErbB2* counts in MCF7 cells are not significantly different between extraction methods. However, *ACTB* counts from ‘bulk’ are significantly higher than from ‘sc’ or ‘cl’ (p < 0.001, Mann-Whitney test with Bonferroni correction). Although, *ACTB* is considered to be a housekeeping gene, its variability due to possible uncontrolled conditions is already described [64]. In this given case we assume, the variability might be related to the different passage numbers (**Tab S4**) or to a different confluency state of the cell culture. Overall, the results of absolute gene mRNA counts of total bulk RNA isolation methods and similarly the results of the crude lysates confirm unbiased, quantitative transcript detection by our scRT-ddPCR method (**Fig 3b**). The larger CVs of single-cell data compared to ‘bulk’ and ‘cl’ (**Tab S5**) are expected and considered to recapitulate expression variability.

**Figure 3:**
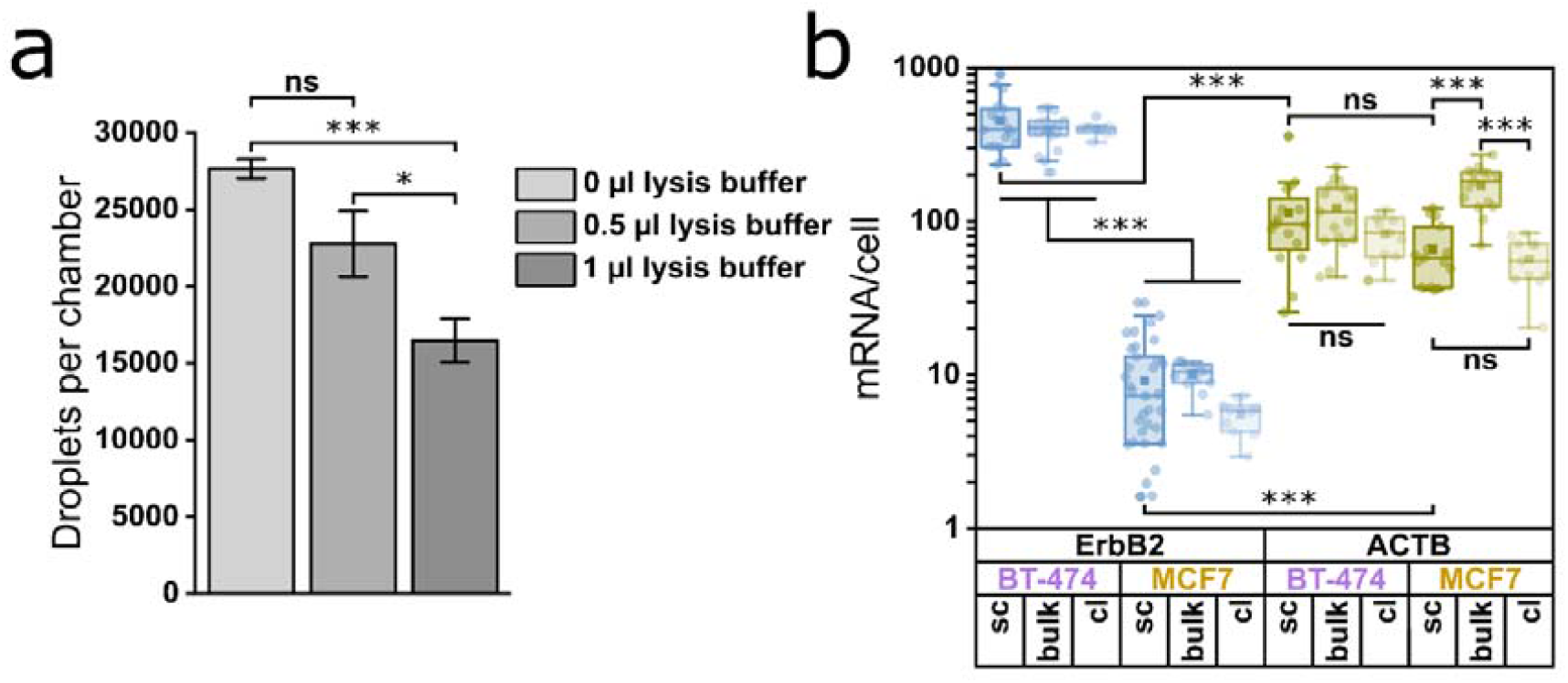
Validation of scRT-ddPCR workflow. **a)** Impact of lysis buffer volume on the number of droplets generated per reaction chamber of one Sapphire Chip. Bar plots show mean values with standard deviation as error bars. Groups were compared using student’s t-test with Bonferroni correction (n = 3). **b)** Absolute gene mRNA per cell counts from different methods (‘sc’ = scRT-ddPCR; ‘bulk’ = quantification from bulk isolated RNA; ‘cl’ = quantification from a crude lysate) according to the genes *ErbB2* and *ACTB* and the cell lines BT-474 and MCF7. Groups were compared using Mann-Whitney test with Bonferroni correction (n ≥ 11). Significance levels not indicated are non-relevant comparisons for this work.

### 3.3 Comparison of down-scaled SMART-Seq® and scRT-ddPCR

Conclusions made solely on the basis of scRNA-seq might be biased because of noise and dropouts and thus need confirmation by PCR means [25]. Because of increased sensitivity, absolute quantification and higher tolerance towards inhibitors [17,32,36,37], we chose scRT-ddPCR for unbiased validation of scRNA-seq data. Furthermore, dPCR provides an orthogonal validation as mRNAs are non-competitively but simultaneously transcribed into cDNAs (partitioning). Thus, the detection events are independent, while in scRNA-seq multiple mRNAs are reverse transcribed in a bulk reaction resulting in competitions and increased propensity for dropouts. Further, we sought to minimize biological and technical variability between the methods by using cells from the same population and the same high precision instrumentation regarding single-cell isolation and liquid handling. We constructed signal distributions of *ErbB2* and *ACTB* expression in MCF7 and BT-474 cells using TPM values from salmon and kallisto, raw counts from STAR aligner or absolute gene mRNA counts per cell from scRT-ddPCR and normalized them to the maximum value per dataset (based on values from **Fig 2b** and **3b**). For *ErbB2* expression in MCF7 cells, we found for scRNA-seq a typical zero-inflation for low abundant targets (∼ 80 % of cells in the first bin; **Fig 4a**, strongly skewed distributions **Tab S6**) [13,14,65]. We observed this behavior also for already published down-scaled Smart-seq2 data from Isakova *et al*. [4]. However, this distribution differs from our scRNA-seq pipelines (Kolmogorov-Smirnov test with Bonferroni multiple testing correction). For the *ErbB2* signal distribution from MCF7 scRT-ddPCR data, we observed a significantly different shape as we could not observe an accumulation of cells in a bin of the histogram and the skewness is much lower (**Tab S6**). For high-abundant transcripts such as *ErbB2* in BT-474 cells, we could detect differences between the alignment tools especially between salmon and kallisto, and STAR. This difference might be justified by the missing normalization of raw counts from STAR aligner or by using the genome as alignment reference. However, *ACTB* signal distributions show no such behavior, but data from Isakova *et al*. are strikingly different compared to all our approaches (**Fig 4b**). Based on the expression values (**Fig 2b** and **Fig 3b**), we calculated log2FCs (MCF7 vs. BT-474) (**Fig 4c**). Additionally, we bootstrapped and down-sampled the scRNA-seq groups to the same sample size as the scRT-ddPCR group to eliminate subsampling errors (**Fig 4c**, shaded bars). The so calculated log2FCs do not differ from the log2FCs calculated by DESeq2 (blue and green arrows; **Fig 2c, 4c** and **Tab S2**). All *ACTB* log2FCs from scRNA-seq and scRT-ddPCR fluctuate within 0 ± 1, which is the null hypothesis of DESeq2 [56], meaning that there is no differential expression between cell lines (**Fig 4c**). The fluctuation probably depicts statistical noise. To compare log2FCs between methods, we bootstrapped their ratio and calculated 95 % CIs. If these 95 % CIs overlap with 1, the methods are assumed to determine the same log2FC (**Eq 1** and **Fig S4**). Log2FCs from both, scRNA-seq (with salmon, kallisto or STAR aligner) and scRT-ddPCR, confirm the overexpression of *ErbB2* in BT-474 cells, although to a significantly different extent, while we could not detect any difference between the log2FCs from the different aligners (**Fig 4c**). scRT-ddPCR predicts significantly stronger overexpression of *ErbB2* in BT-474 cells. This can potentially be explained by the biased detection of *ErbB2* expression in MCF7 cells by scRNA-seq (**Fig 4a**).

**Figure 4:**
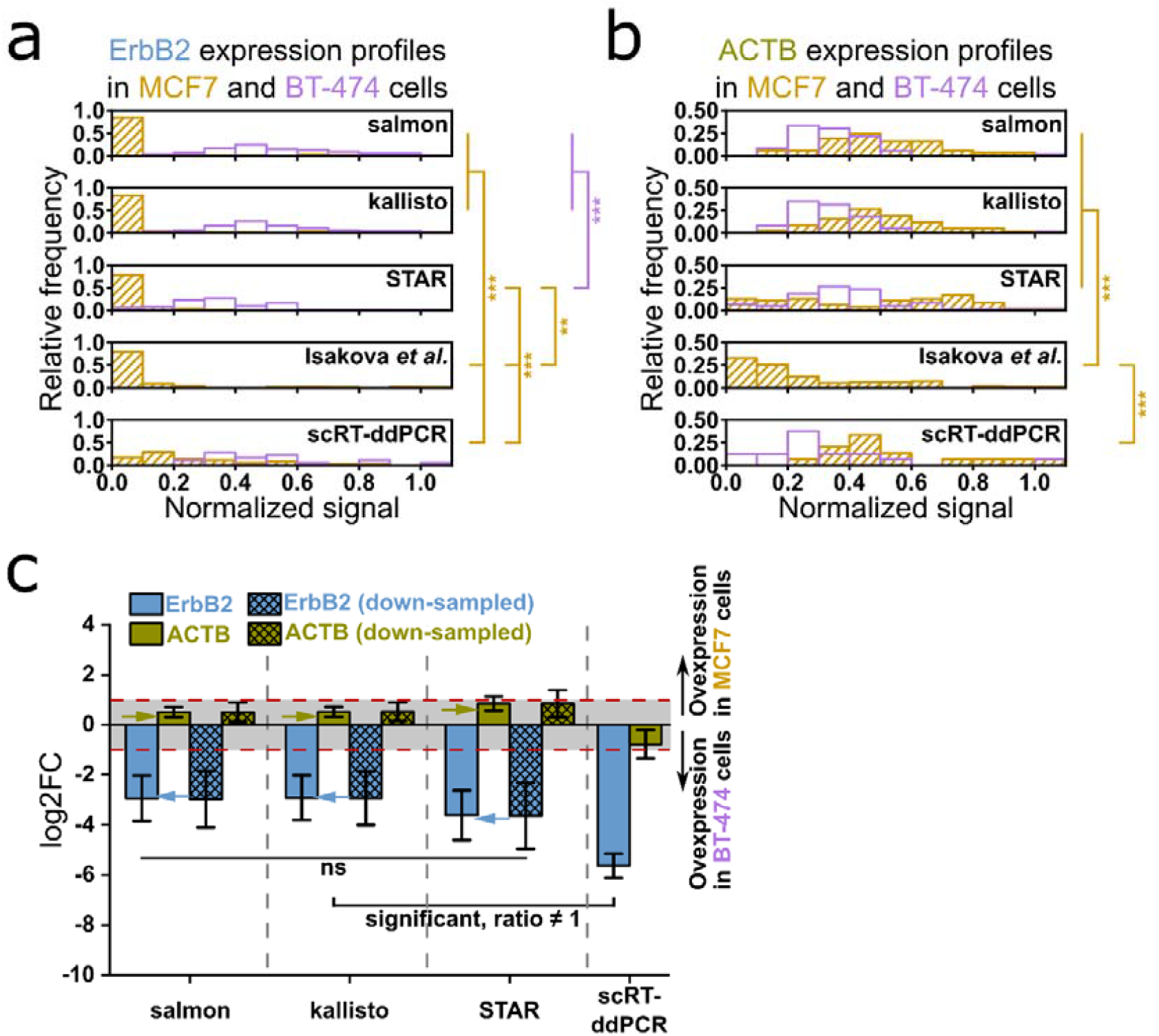
Comparison of scRT-ddPCR and scRNA-seq on the basis of signal distributions and fold changes. **a)** Distribution of normalized *ErbB2* expression signal (normalized to maximum signal) from salmon, kallisto and STAR aligner used in this study, MCF7 expression data from Isakova *et al*. (10-fold down-scaled Smart-seq2) and scRT-ddPCR in MCF7 and BT-474 cells. All distributions were compared using a Kolmogorov-Smirnov test with Bonferroni correction for multiple testing. Non-significant difference are not shown. **b)** Distribution of normalized *ACTB* expression signal (normalized to maximum signal) from salmon, kallisto and STAR aligner used in this study, MCF7 expression data from Isakova *et al*. (1/10 down-scaled Smart-seq2) and scRT-ddPCR in MCF7 and BT-474 cells. All distributions were compared using a Kolmogorov-Smirnov test with Bonferroni correction for multiple testing. Non-significant difference are not shown. **c)** Log2FCs (MCF7 vs. BT-474) for scRNA-seq data processed with salmon, kallisto and STAR aligner and scRT-ddPCR, calculated on the basis of expression values as shown in **Fig 2b**. The shaded bars show log2FCs of the respectives group down-sampled by bootstrapping to the same sample size of scRT-ddPCR. Log2FCs were compared using a bootstrapping comparison (2.8 Bootstrapping comparison and **Fig S4**). Bars depict mean log2FCs with bootstrapped error bars indicating 95 % CI. Log2FCs within 0 ± 1 are not considered to be statistically significant. Blue and green arrows indicate log2FCs calculated by DESeq2.

## 4 Discussion

In this study, we present a novel, orthogonal method, scRT-ddPCR, for the validation of scRNA-seq fold changes. Durst *et al*. found that absolute quantification is the most reliable approach for single-cell analysis [39], which is the key feature of dPCR and delivers a ground truth that facilitates inter-experimental comparisons as it is detached from any standard. This is achieved by the inherent partitioning of dPCR, which further allows spatially separated but simultaneous reverse transcription of mRNAs. This potentially improves cDNA capture through enrichment. Thus, for low-abundant transcripts, which are often referred to as highly interesting but difficult to reliably analyze [15–18], dPCR might therefore be of great advantage.

First, we aimed to enhance molecular detections for SMART-Seq® by down-scaling, which is frequently applied to scRNA-seq protocols to increase throughput, reduce costs, and increase sensitivity, while maintaining data quality [4,59–62]. We demonstrated that our down-scaled SMART-Seq® protocol using F.SIGHT™ and I.DOT delivers high quality data (**Fig 1, S1** and **Tab S1**). We validated our method by comparative UMAP-clustering against published, down-scalded data [4] and found excellent conformity (**Fig 2a**). Compared to 3’-counting methods such as the Chromium system [24], full-length protocols such as SMART-Seq® have already demonstrated better coverage of low-abundant transcripts [3,23], but are still not sensitive enough to detect low-abundant *ErbB2* mRNA in MCF7 cells as we show (**Fig 4a**). We relate these dropouts to the simultaneous reverse transcription of multiple poly(A)-mRNAs into cDNAs. Thus, our data support previous findings of dropouts in scRNA-seq [13,14].

Secondly, we successfully validated our scRT-ddPCR method (**Fig 3** and **S2**), which underlines that scRT-ddPCR can serve as a ground truth. We could detect *ErbB2* expression in MCF7 cells without dropouts (**Fig 4a**) potentially because of the inherent partitioning step in dPCR, which increases the effective mRNA concentration [36]. *ErbB2* expression in BT-474 cells and *ACTB* expression in MCF7 and BT-474 cells could be similarly detected (**Fig 3b, 4a** and **4b**). Thus, our proposed scRT-ddPCR method can reliably and absolutely quantify low- and high-abundant transcripts offering a solution for fold change validation. However, a drawback of dPCRs is the degree of multiplexing, which limits the genes to analyze. At the time we conducted the experiments, the highest degree of multiplexing was three colors [46]. Recent developments in dPCR instrumentation allow five (QIAcuity, Qiagen) or six (Prims6, Stilla Technlogies) color detection. Most of the existing approaches for scRNA-seq validation use qPCR [26–30], but also these cyclers do not offer higher degree of multiplexing than six. Alternatively, approaches of monochrome multiplexing, such as photo bleaching, could be used to extend the degree of multiplexing beyond hardware limitations [66,67].

Finally, we compared log2FCs from scRNA-seq and scRT-ddPCR and found that *ACTB* log2FCs from scRNA-seq were not different from scRT-ddPCR log2FCs (**Fig 4c**). In both cell lines, *ACTB* expression has a good signal distribution for scRT-ddPCR and all aligners used in scRNA-seq (**Fig 4b** and **Tab S6**). Strikingly, the signal distribution obtained from published data shows a much stronger skewness (**Fig 4b** and **Tab S6**), which could be an indication that our down-scaled protocol using F.SIGHT™ and I.DOT performs better than existing down-scaled versions. While *ErbB2* fold changes are consistent among the aligners used, we found a significant difference between scRNA-seq and scRT-ddPCR (**Fig 4c**). We hypothesize that these differences originate from the heavily skewed signal distributions (skewness ≈ 3) of *ErbB2* in MCF7 cells, which indicate dropouts (**Fig 4a** and **Tab S6**). However, scRNA-seq and scRT-ddPCR use different priming strategies for cDNA synthesis [68] and in scRNA-seq protocols more PCR steps are included (i.e. cDNA amplification, tagmented library amplification, bridge amplification), which potentiate biases and promote dropouts. Biases in scRNA-seq could also originate from the bioinformatics tools used as we observe that some genes are predicted to be overexpressed with some aligners (**Fig 2c** and **2d**) and that overexpression is not consistent across the aligners (**Tab S2**).

In this study, we only evaluate the impact of dropouts on individual fold changes but we assume that this has far-reaching implications on DE analyses and their conclusions, especially since this is not an issue limited to scRNA-seq but also exists in conventional RNA-seq [69–71]. However, it is pronounce in scRNA-seq because of low sample input [13,14]. In concordance with this, we could show that the alignment tool has an impact on the amount of DEGs and on the fold changes (**Fig 2c, 2d, 4c** and **Tab S2**). This underlines the necessity for an independent validation method that allows the reliable detection of absolute mRNA counts such as our scRT-ddPCR. Our here presented scRT-ddPCR method can thus serve as a platform for mRNA analysis but could also be extended to the protein [31] and DNA analysis [72] or different cell types [45]. On top of that, it is compatible with any plate-based sequencing protocol such as Smart-seq3, Smart-seq3xpress or FLASH-seq [60,61,73]. In conclusion, we think this method is a valuable addition to the toolbox of researchers interested in single-cell transcriptomics because of its reliability, ease of use, reduced costs, and increased sensitivity.

## Abbreviations

CI: confidence interval,
cl: crude lysate,
CV: coefficient of variance,
DE: differential expression,
DEG: differentially expressed gene,
dPCR: digital PCR,
FACS: fluorescence-activated cell sorting,
log2FC: log2 of fold change between two conditions,
RT-qPCR: reverse transcription quantitative PCR,
sc: single-cell,
scRNA-seq: single-cell RNA sequencing,
scRT-ddPCR: single-cell reverse transcription droplet digital PCR,
TPM: Transcripts per kilobase million,
UMAP: uniform manifold approximation and projection,

## Author contribution statement

Conceptualization: Csaba Jeney, Tobias Lange

Investigation: Tobias Lange, Tobias Groß

Methodology: Tobias Lange, Julia Scherzinger, Elly Sinkala, Ábris Jeney, Csaba Jeney, Tobias Groß

Formal analysis: Tobias Lange, Ábris Jeney

Supervision and funding acquisition: Csaba Jeney, Stefan Zimmermann, Peter Koltay, Felix von Stetten, Roland Zengerle

Writing – original draft: Tobias Lange

Writing – review and editing: Csaba Jeney, Stefan Zimmermann, Peter Koltay, Felix von Stetten, Roland Zengerle, Christoph Niemöller, Tobias Lange, Tobias Groß, Ábris Jeney

All authors read and approved the final version of the manuscript.

## Declaration of competing interest

J.S., E.S. and C.N. are employees of CYTENA GmbH, which produces the F.SIGHT™ single-cell dispenser used in this study. T.L., T.G., P.K, and C.J. are employees of Actome GmbH, which develops the LBT lysis buffer used in this study and P.K., R.Z. and C.J. are shareholders of Actome GmbH. The remaining authors declare no competing interest.

## Data and code availability

scRNA-seq data is available from GEO under accession number: GSE201443. Jupyter notebooks for the analysis of our scRNA-seq data are available from github.com/LangeTo/scRNA-seq_scripts. The code for bootstrapping comparison is available upon reasonable request.

## Acknowledgments

We would like to thank the Actome GmbH for providing research licenses, for help with laboratory work and for advices on the manuscript. We would like to acknowledge the Lighthouse Core Facility of the Universitätsklinkium Freiburg for assistance with dPCR. Lighthouse Core Facility is funded in part by the Medical Faculty, University of Freiburg (Project Number 2021/A2-Fol). We appreciate the RNA isolation kit gift form Zymo Research Europe (Freiburg). We acknowledge Martin Adam Stoffel (University of Edinburgh) for ideas on bootstrapping. This work was supported by grants from the Baden-Württemberg Stiftung in the framework of “Methoden in den Lebenswissenschaften” (MONOGRAM – MET-ID55) and the Ministerium für Wissenschaft, Forschung und Kunst Baden-Württemberg (DINAMIK – 7533-7-11.10-6).

---

**Graphical abstract.**
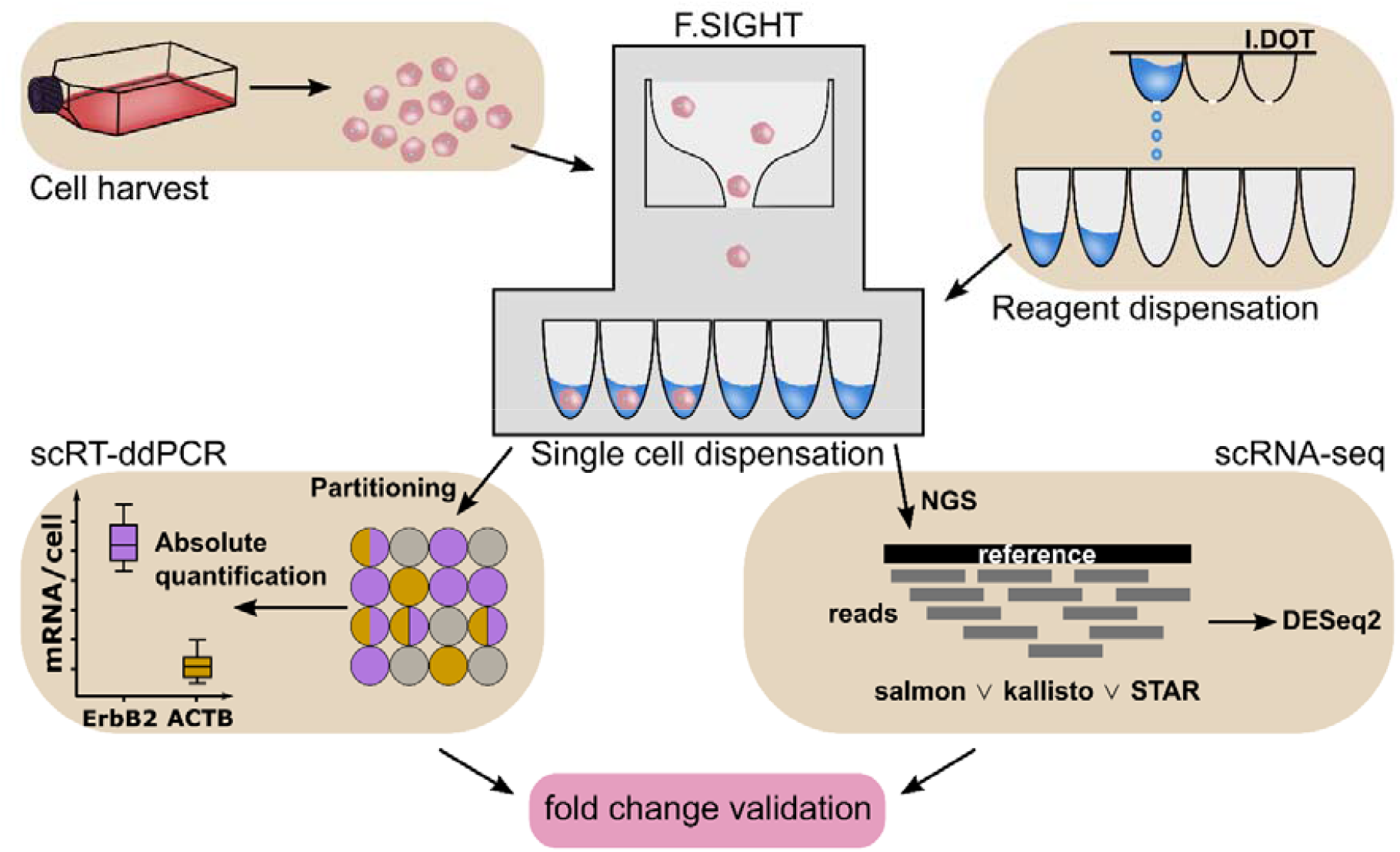
Validation of scRNA-seq fold changes by scRT-ddPCR.

## Notes

### Summary of Updates

Please check the TRiP Peer Reviews tab for detailed information on the revisions of this manuscript.

