## Supplementary figures S1 to S4 for "Validation of scRNA-seq by scRT-ddPCR using the example of *ErbB2* in MCF7 cells"

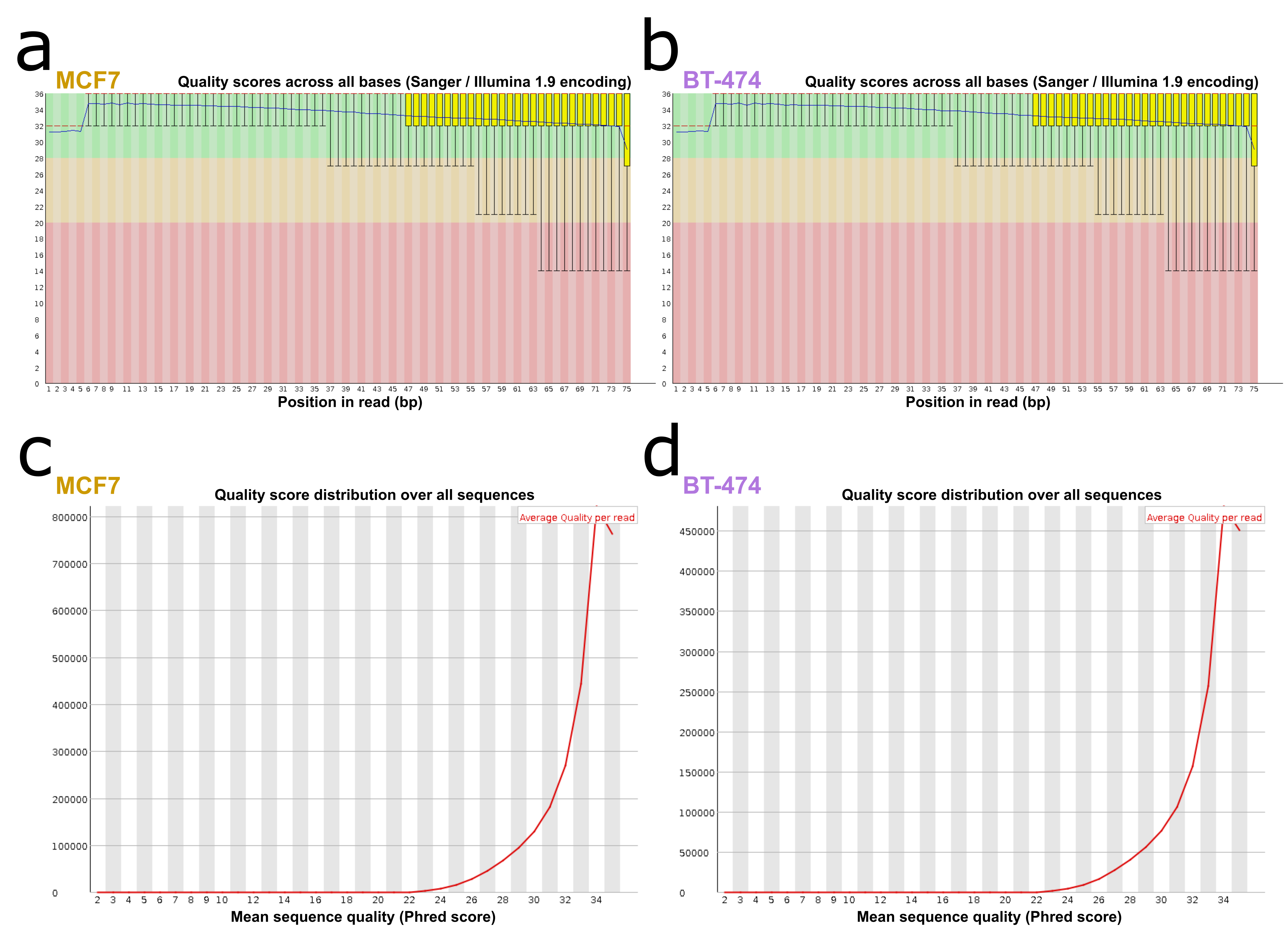

**Figure S1: scRNA-seq quality control by FastQC. a, b)** Per base sequence quality of one representative MCF7 and BT-474 cell. **c, d)** Per sequence quality scores of one representative MCF7 and BT-474 cell.

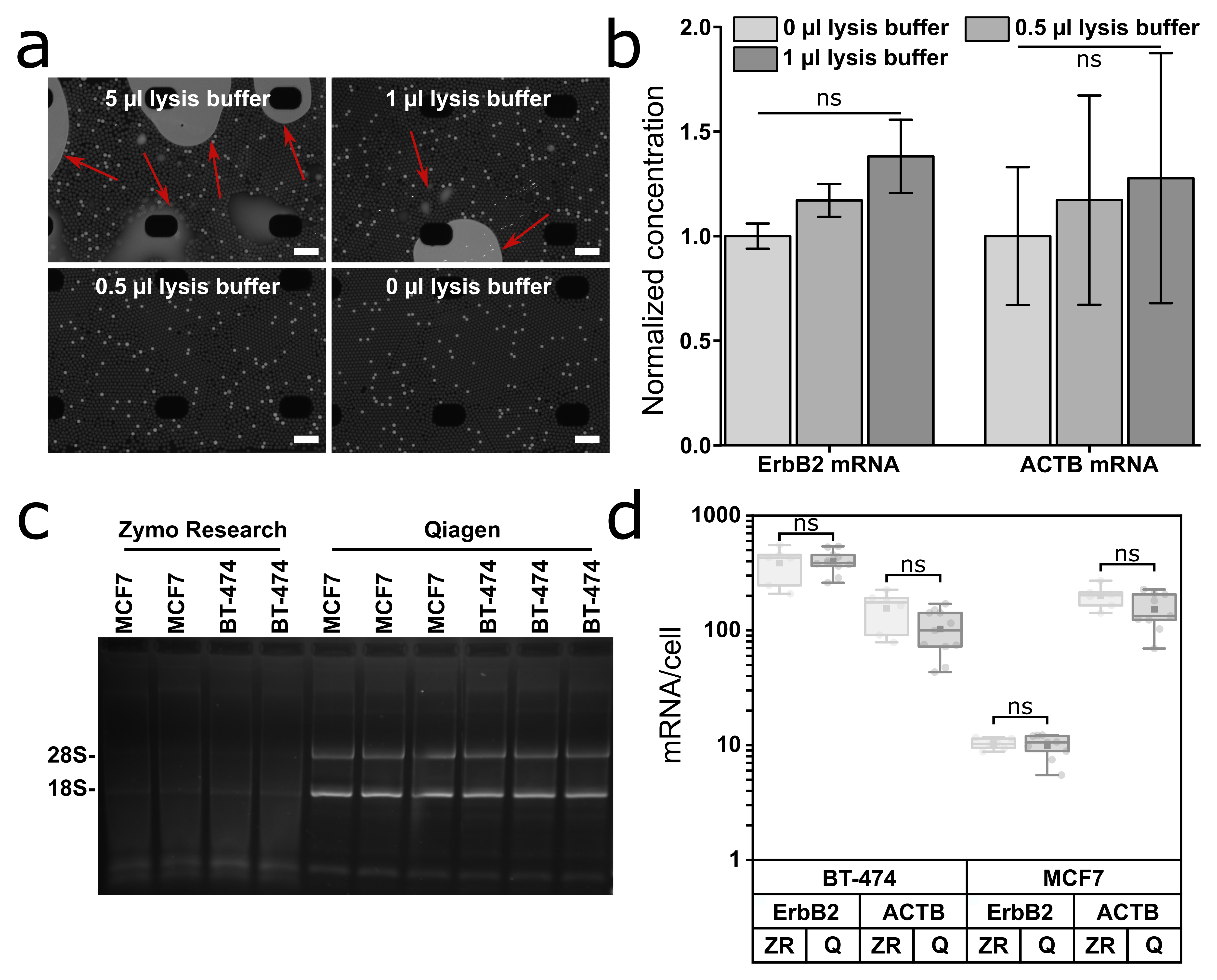

**Figure S2: Impact of lysis buffer volume on droplet generation in RT-ddPCR and impact of different RNA isolation methods on gene mRNA per cell yield. a)** 5 µl, 1 µl, 0.5 µl lysis buffer were spiked into a RT-ddPCR. The raw images from the Crystal Miner software are shown. Black areas are support structures. Areas of coalescence are highlighted with red arrows. Scale bars: 664 µm. **b)** Absolute mRNA concentrations of *ErbB2* and *ACTB* in the presence of 0.5 µl and 1 µl lysis buffer were determined and normalized to 0 µl lysis buffer. All groups per gene were compared using Student’s t-test with Bonferroni correction (n = 3). **c)** Native agarose gel of total isolated RNA by kits from Zymo Research and Qiagen to access the quality of RNA. **d)** Comparison of gene mRNA per cell counts for *ErbB2* and *ACTB* in MCF7 and BT-474 cell lines using kits from different manufacturers (ZR: Zymo Research using mechanic lysate homogenization; Q: Qiagen using enzymatic lysate homogenization). ZR and Q groups were compared using Student’s t-test (n ≥ 6). Data was subsumed for further comparisons.

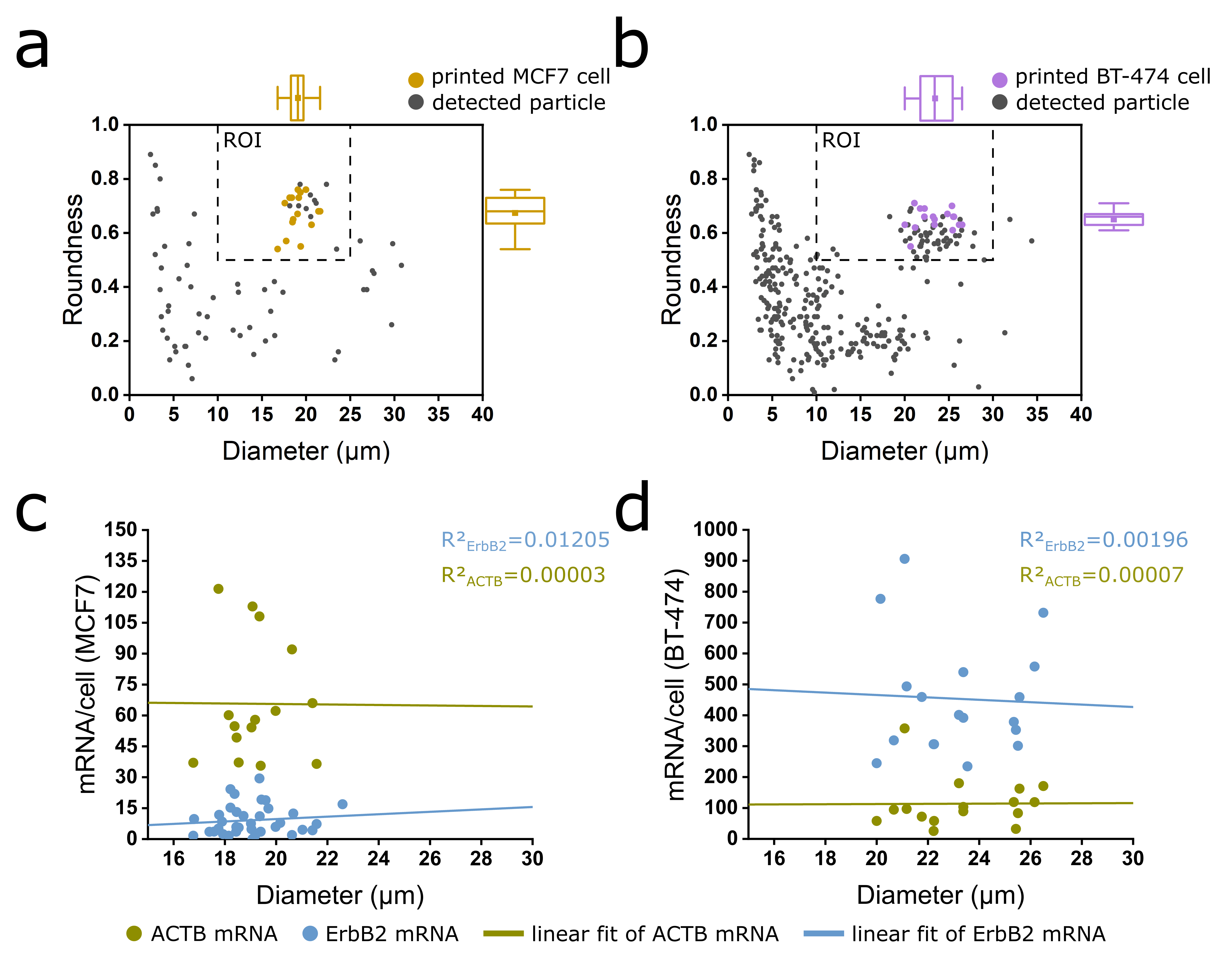

**Figure S3: Documentation of single-cell dispensation process and correlation of gene mRNA per cell counts with cell size from scRT-ddPCR. a, b)** Scatter plots (roundness vs. diameter) of printed MCF7 and BT-474 cells (colored dots) or detected particles (grey dots). Boxplots show roundness and diameter distributions of printed cells. ROI (region of interest) depicts the desired morphological criteria. **c, d)** Correlation of gene mRNA per cell counts with the cell diameter. Pearson’s correlation coefficient is indicated.

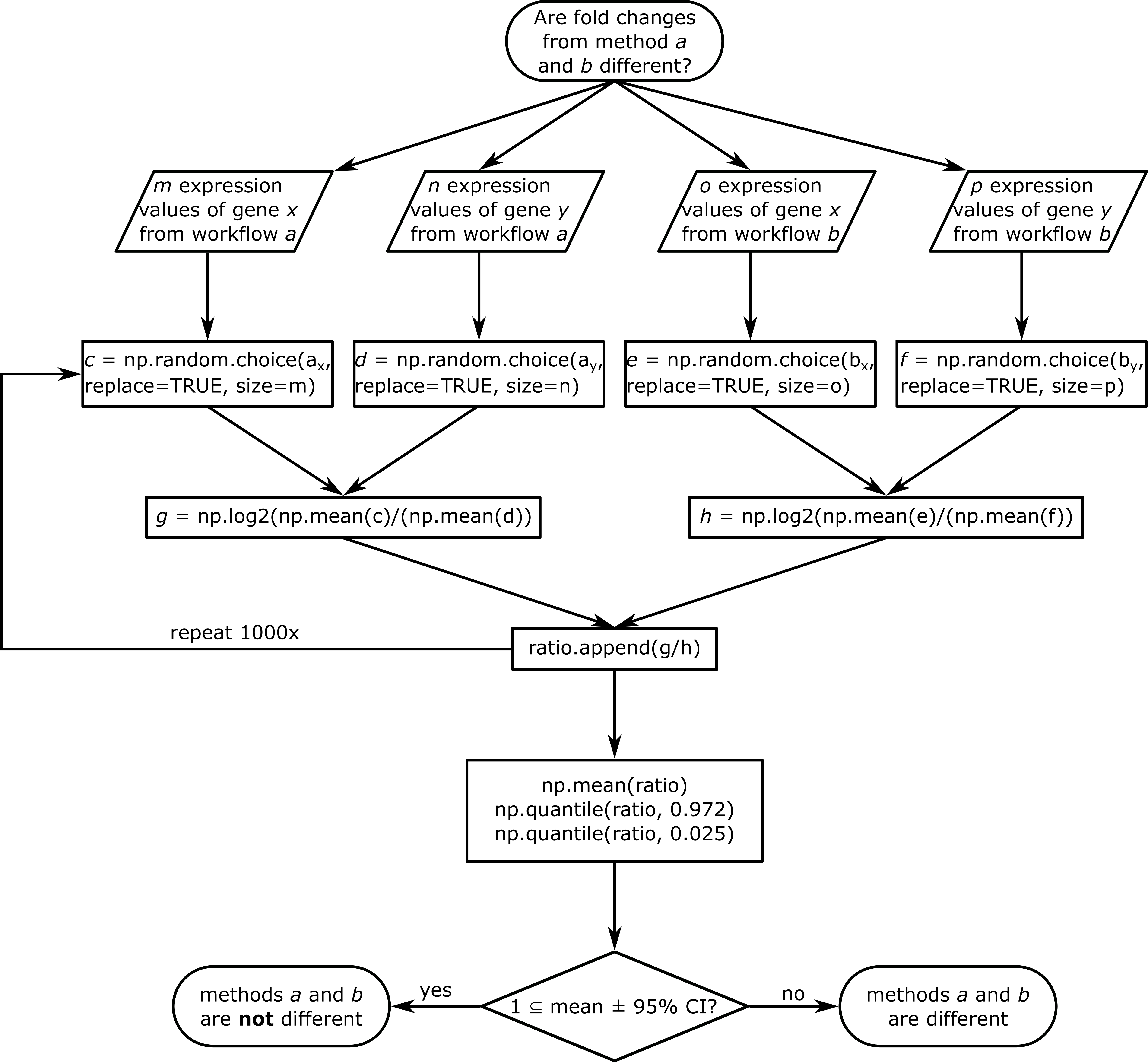

**Figure S4: Schematic of bootstrapping comparison algorithm.** The algorithm needs four input arrays containing arbitrary amounts of expression values from two genes and two methods. In the next step, random subsamples with the same length as the original input arrays are drawn with replacement using numpy’s np.random.choice() command. The mean value of each subsample is calculated and the log2FCs are determined according to **Eq 1**. The ratio of these two fold changes is appended to the new array ‘ratio’. This calculation is performed 1,000 times. Ultimately, the mean of the array ‘ratio’ and its 95 % CIs are calculated. If 1 is a subset of the mean and its 95 % CI, the log2FCs of the two methods are not significantly different.
