## Supplementary tables S1 to S7 for "Validation of scRNA-seq by scRT-ddPCR using the example of *ErbB2* in MCF7 cells"

**Table S1**: Filtering of cells for scRNA-seq downstream anylsis as shown by the amount (fraction) of cells excluded by different quality control steps. Inclusion criteria: single-cell isolation analysis: single cell encapsulation manually verified by image-based analysis (**Fig 1a**); alignment quality: salmon and kallisto: alignment efficiency ≥ 80 % (**Fig 1d**), STAR: fraction of uniquely mapped reads ≥ 60 % (**Fig 1e**); transcript count ≥ 1E+5 (**Fig 1f**).

| **Cell line** | **Initial cell amount** | **Single-cell isolation analysis** | **Alignment quality** | | | **Transcript count** | | |
| --- | --- | --- | --- | --- | --- | --- | --- | --- |
|  |  |  | salmon | kallisto | STAR | salmon | kallisto | STAR^§^ |
| **MCF7** | 84 | 6^$^ | 20 (26 %) | 20 (26 %) | 29 (37 %) | 1  (2 %) | 1  (2 %) | 2  (4 %) |
|  | **Number of MCF7 cells in downstream analysis** | | | | | 57 | 57 | 47 |
| **BT-474** | 84 | 3^#^ | 19 (23 %) | 17 (21 %) | 14 (17 %) | 2  (3 %) | 4  (6 %) | 7 (10 %) |
|  | **Number of BT-474 cells in downstream analysis** | | | | | 60 | 60 | 60 |

^$^Corresponding to a single-cell isolation efficiency of 93 %.

^#^Corresponding to a single-cell isolation efficiency of 96 %.

^§^The average fraction of uniquely mapped reads after filtering was 84.7 % for MCF7 cells and 83.5 % for BT‑474 cells.

**Table S2**: Expression values from genes of interest *ErbB2*, *ACTB*, *KRT8* and *TFF1* after DESeq2 analysis with input from different aligners (see **Fig 2c**).

| **Aligner** | **Gene** | **baseMean** | **log2FC** | **Adjusted p-value** | **Overexpression │log2FC│ > 1 and adjusted p-value < 0.05** |
| --- | --- | --- | --- | --- | --- |
| **salmon** | *ErbB2* | 361.87971 | -2.8578504 | 7.23E-16 | BT-474 |
|  | *ACTB* | 25993.6337 | 0.33248271 | 4.91E-04 | - |
|  | *KRT8* | 10557.9592 | 0.80857214 | 2.52E-11 | - |
|  | *TFF1* | 8934.60496 | 1.88158285 | 1.08E-19 | MCF7 |
| **kallisto** | *ErbB2* | 342.081257 | -2.88952331 | 1.51E-17 | BT-474 |
|  | *ACTB* | 25819.8298 | 0.33421852 | 3.77E-04 | - |
|  | *KRT8* | 9005.88508 | -0.31790799 | 1.38E-01 | - |
|  | *TFF1* | 9167.33808 | 1.90108326 | 4.77E-20 | MCF7 |
| **STAR** | *ErbB2* | 181.654883 | -3.7519021 | 2.31E-56 | BT-474 |
|  | *ACTB* | 14690.6213 | 0.60709742 | 5.86E-13 | - |
|  | *KRT8* | 4947.28588 | 1.28277102 | 5.36E-94 | MCF7 |
|  | *TFF1* | 5434.42135 | 2.06371347 | 4.39E-94 | MCF7 |

**Table S3**: Total RNA concentration and RNA purity after bulk RNA isolation from 1x10^6^ MCF7 cells and 1x10^6^ BT-474 cells using Zymo Research (enzymatic lysate homogenization) or Qiagen kits (mechanic lysate homogenization).

| **Isolation Procedure** | **Cell line** | **Concentration [ng/µl]** | **Total amount [µg]** | **A260/A280** | **A260/A230** |
| --- | --- | --- | --- | --- | --- |
| **Zymo Research** | **MCF7** | 1972.8 | 29.6 | 2.15 | 2.22 |
|  |  | 1755.1 | 26.3 | 2.13 | 2.20 |
|  | **BT-474** | 1178.0 | 17.7 | 2.11 | 2.16 |
|  |  | 1381.5 | 20.7 | 2.11 | 2.19 |
| **Qiagen** | **MCF7** | 743.0 | 22.3 | 2.16 | 1.46 |
|  |  | 779.2 | 23.4 | 2.15 | 1.97 |
|  |  | 741.5 | 22.2 | 2.14 | 2.24 |
|  | **BT-474** | 523.1 | 15.7 | 2.15 | 2.17 |
|  |  | 499.2 | 15.0 | 2.15 | 1.32 |
|  |  | 526.6 | 15.8 | 2.16 | 2.14 |

**Table S4**: Passage numbers of cells used for the indicated methods.

| **Cell Line** | **scRNA-seq** | **RT-ddPCR** | | |
| --- | --- | --- | --- | --- |
|  |  | **sc** | **bulk** | **cl** |
| **MCF7** | 10 | 10 | 11 | 10 |
| **BT-474** | 13 | 13 | 14 | 14 |

**Table S5**: CVs of gene mRNA per cell counts from different methods for MCF7 and BT-474 cells as well as for the genes *ErbB2* and *ACTB*.

| **Cell Line** | **Gene** | **RT-ddPCR** | | |
| --- | --- | --- | --- | --- |
|  |  | **sc** | **bulk** | **cl** |
| **MCF7** | ***ErbB2*** | 78.8 % | 18.6 % | 24.5 % |
| **MCF7** | ***ACTB*** | 44.3 % | 31.7 % | 35.1 % |
| **BT-474** | ***ErbB2*** | 41.6 % | 25.5 % | 9.6 % |
| **BT-474** | ***ACTB*** | 69.6 % | 44.7 % | 32.0 % |

**Table S6**: Skewness and kurtosis of signal distributions from **Fig 4a** and **4b**. The skewness of a normal distribution is 0 and its kurtosis 3.

| **Gene** | **Signal distribution** | **Cell line** | **Skewness** | **Kurtosis** |
| --- | --- | --- | --- | --- |
| *ErbB2* | salmon | MCF7 | 3.05 | 9.39 |
|  |  | BT-474 | 0.50 | -0.13 |
|  | kallisto | MCF7 | 2.97 | 8.75 |
|  |  | BT-474 | 0.58 | -7.52E-4 |
|  | STAR | MCF7 | 3.16 | 9.95 |
|  |  | BT-474 | 0.91 | 1.51 |
|  | Isakova *et al.* | MCF7 | 3.19 | 9.79 |
|  | scRT-ddPCR | MCF7 | 1.07 | 0.62 |
|  |  | BT-474 | 1.15 | 0.71 |
| *ACTB* | salmon | MCF7 | 0.48 | 0.06 |
|  |  | BT-474 | 2.13 | 9.17 |
|  | kallisto | MCF7 | 0.44 | 0.11 |
|  |  | BT-474 | 2.29 | 10.36 |
|  | STAR | MCF7 | -0.12 | -1.33 |
|  |  | BT-474 | 0.72 | 1.21 |
|  | Isakova *et al.* | MCF7 | 1.28 | 0.98 |
|  | scRT-ddPCR | MCF7 | 0.89 | -0.49 |
|  |  | BT-474 | 2.04 | 5.61 |

**Table S7**: Compliance with dMIQE guidelines for essential information.

| **Item** | **Comment** |
| --- | --- |
| **Experimental design** |  |
| Definition of experimental and control groups | ‘sc’ (scRT-ddPCR); controls: ‘bulk’ (quantification from isolated RNA from bulk cells), ‘cl’ (quantification from a crude lysate of bulk cells) |
| Number of single cells within each group | \| MCF7 \| \| BT-474 \|; \| *ErbB2* \| *ACTB* \| *ErbB2* \| *ACTB*; ‘sc’ \| 35 \| 15 \| 18 \| 16; ‘bulk’ \| 15 \| 15 \| 17 \| 17; ‘cl’ \| 11 \| 11 \| 11 \| 11 |
| **Sample** |  |
| Volume of mass of sample processed | Single cells |
| Microdissection or macrodissection | Not applicable |
| Processing procedure | ‘sc’: single cells dispensed into lysis buffer, ‘bulk’: serial dilution of total isolated RNA, ‘cl’: single-cell volume equivalent dispensed into dPCR master mix |
| If frozen – how and how quickly? | Not frozen |
| If fixed – with what, how quickly? | Not fixed |
| Sample storage conditions and duration (especially for formalin-fixed, paraffin-embedded samples) | Not applicable |
| **Nucleic acid extraction** |  |
| Quantification – instrument/method | NanoDrop™ One |
| Storage conditions: temperature, concentration, duration, buffer | ‘sc’ and ‘cl’: direct processing, ‘bulk’ ‑20 °C, 500 to 2000 ng/µl, 1 week, H_2_O |
| DNA or RNA quantification | RNA |
| Quality/integrity, instrument/method | Gel electrophoresis (**Fig S2c**) |
| Template structural information | Not provided |
| Template modification | None |
| Template treatment (initial heating or chemical denaturation) | None |
| Inhibition dilution or spike | None |
| DNA contamination assessment of RNA sample | Gel electrophoresis (**Fig S2c**) |
| Details of DNase treatment where performed | On-column DNase I digest in Zymo Research RNA isolation protocol (2.2 Total RNA isolation and bulk cell lysis) |
| Storage of nucleic acid: temperature, concentration, duration, buffer | ‘sc’ and ‘cl’: direct processing, ‘bulk’*: ‑*20 °C, 500 to 2000 ng/µl, 1 week, H_2_O |
| **RT** |  |
| cDNA priming method + concentration | Gene-specific, 900 nM |
| One- or 2-step protocol | One-step protocol |
| Amount of RNA used per reaction | ‘sc’: single cells, ‘cl’: single-cell volume equivalent, ‘bulk’: serial dilution between 0.5 ng and 40 ng |
| Detailed reaction components and conditions | 2.5 Droplet digital PCR |
| **dPCR target information** |  |
| Sequence accession number | *ErbB2*: NM_001005862.2, *ACTB*: NM_001101.3 |
| Amplicon length | *ErbB2*: 60 bp, *ACTB*: 63 bp |
| *In-silico* specificity screen | Not applicable because commercial assay used |
| Location of each primer by exon or intron | Both assays are exon spanning: *ErbB2*: exon boundary 17 – 18, *ACTB*: exon boundary 2 – 3 |
| Where appropriate, which splice variants are targeted? | No splice variants targeted |
| **dPCR oligonucleotides** |  |
| Primer sequences and/or amplicon context sequence | Amplicon sequence: *ErbB2*: GGAGGCTGACCAGTGTGTGGCCTGTGCCCACTATAAGGAC CCTCCCTTCTGCGTGGCCCG, *ACTB*: GGCGTGATGGTGGGCATGGGTCAGAAGGATTCCTATGTGGG CGACGAGGCCCAGAGCAAGAGA |
| Location and identity of any modifications | No modifications |
| **dPCR protocol** |  |
| Reaction volume and amount of RNA/cDNA/DNA | ‘sc’: single cells, ‘cl’: single-cell volume equivalent (100 nl), ‘bulk’: 1 µl of serial dilution of total isolated RNA |
| Primer, probe, Mg^2+^, and dNTP concentrations | Primer: 900 nM, probe: 250 nM, Mg^2+^: not spefied, dNTP: not specified |
| Polymerase identity and concentration | Not specified |
| Buffer/kit catalogue no. and manufacturer | 2.5 Droplet digital PCR |
| Additives | Magnesium chloride, RNase inhibitor protein, AccuVue blue qPCR dye, stabilizers |
| Complete thermocycling parameters | 2.5 Droplet digital PCR |
| Partition number | 25345 (20 %CV) partitions on average across all experiments |
| Individual partition volume | 0.548 nl |
| Total volume of the partitions measured (effective reaction size) | 13.9 µl on average |
| Comprehensive details and appropriate use of controls | NTC (‘sc’: no cell dispensed, ‘cl’: PBS dispensed, ‘bulk’: 1 µl of PBS added) |
| Manufacturer of dPCR instrument | Stilla Technologies |
| **dPCR validation** |  |
| Specificity (when measuring rare mutations, pathogen sequences etc.) | Not applicable |
| If multiplexing, comparison with singleplex assays | Not performed |
| **Data analysis** |  |
| Mean copies per partitions | λ = 0.71 (267 % CV) (overall average) |
| dPCR analysis program | Crystal Miner™ |
| Outlier identification and disposition | Not applicable |
| Results of no-template controls | All negative (data not shown) |
| Examples of positive and negative experimental results as supplemental data | **Fig S2a** |
| Where appropriate, justification of number and choice of reference genes | Not applicable |
| Where appropriate, description of normalization method | Not applicable |
| Number and stage of technical replicates | 2 to 3 |
| Repeatability | Proven with different RNA isolation kits (**Fig S2d**) |
| Experimental variance or CI | **Fig 3b** and **Table S5** |
| Statistical methods used for analysis | 2.7 Statistical analysis and 2.8 Bootstrapping fold change ratio comparison |
